# In Vitro Activities of Daptomycin Combined with Fosfomycin or Rifampin on Planktonic and Adherent Linezolid-resistant *Enterococcus faecalis*

**DOI:** 10.1101/345892

**Authors:** Jin-xin Zheng, Xiang Sun, Zhi-wei Lin, Guo-bin Qi, Hao-peng Tu, Yang Wu, Si-bo Jiang, Zhong Chen, Qi-wen Deng, Di Qu, Zhi jian Yu

**Affiliations:** Department of Infectious Diseases and the Key Lab of Endogenous Infection, Shenzhen Nanshan People’s Hospital and The 6th Affiliated Hospital of Shenzhen University Health Science Center, Shenzhen 518052, China.; Key Laboratory of Medical Molecular Virology of Ministries of Education and Health, School of Basic Medical Science and Institutes of Biomedical Sciences, Shanghai Medical College of Fudan University, Shanghai 200032, China.; Department of Pharmaceutics, University of Florida, Orlando 32827, USA

**Keywords:** *Enterococcus faecalis*, linezolid-resistant, biofilm, daptomycin, fosfomycin, rifampin

## Abstract

This study aimed to explore daptomycin combined with fosfomycin or rifampin against the planktonic and adherent linezolid-resistant isolates of *Enterococcus faecalis*. Four linezolid-resistant isolates of *E. faecalis* which formed biofilms were collected for this study. Biofilm biomasses were detected by crystal violet staining. The adherent cells in the mature biofilms were counted by CFU numbers and observed by confocal laser scanning microscope (CLSM). In time-killing studies, daptomycin combined with fosfomycin or rifampin (4xMIC) demonstrated bactericidal activities on the planktonic cells, and daptomycin combined with fosfomycin killed more planktonic cells (at least 2-log10 CFU/ml) than daptomycin or fosfomycin alone. Daptomycin alone showed activities against the mature biofilms, and daptomycin combined with fosfomycin (16xMIC) demonstrated significantly more activity than daptomycin or fosfomycin alone against the mature biofilms in three of the four isolates. Daptomycin alone effectively killed the adherent cells, and daptomycin combined with fosfomycin (16xMIC) killed more adherent cells than daptomycin or fosfomycin alone in these mature biofilms. The high concentrations of daptomycin (512 mg/L) combined with fosfomycin indicated more activity than 16xMIC of daptomycin combined with fosfomycin on the adherent cells and the mature biofilms. The addition of rifampin increased the activity of daptomycin against the biofilms and the adherent cells of FB-14 and FB-80 isolates, but was not observed in FB-1 and FB-2 isolates. In conclusion, daptomycin combined with fosfomycin works effectively against the planktonic and adherent linezolid-resistant isolates of *E. faecalis*. The role of rifampin in these linezolid-resistant isolates is discrepant and needs more studies.

---

*Enterococcus faecalis* has become one of the most common pathogens of nosocomial infections in the last two decades, which usually causes urinary tract, respiratory tract, peritoneum, and bloodstream infections (1). Of particular concern is the increasing difficult treatment of *E. faecalis* as it has intrinsic and acquired resistance to many antimicrobial agents. The outbreaks of vancomycin-resistant enterococci (VRE) infections attracted global attention due to their extensive resistance to a plethora of antibiotics in recent years (2). Linezolid, was the first antimicrobial agent of the oxazolidinones drugs to treat VRE infections, but the growing cases of linezolid-resistant enterococci have emerged in hospitals with its wide use (3).

In addition to the drug resistance problem, *E. faecalis* has been found with a high capacity for biofilm formation, which makes infections more difficulty to treat. In Britain, 100% of *E. faecalis* strains from bloodstream infections had the ability to form biofilms, and another study from Japan indicated that all of 352 *E. faecalis* isolates in urinary tract infections formed biofilms (4, 5). Other studies also showed that about 50-90% of *E. faecalis* clinical isolates formed biofilms (6-8). Recently, among 265 *E. faecalis* strains from China, 90% of linezolid-resistant isolates were found with different levels of biofilm formation (9).

Biofilms are enclosed within an exopolymer matrix that can restrict the diffusion and penetration of antimicrobials, and thus make them very difficult to erase (10). At present, only a few antimicrobials showed activities on enterococci biofilms. Previous research found that the daptomycin has more activity than linezolid against biofilm-forming *E. faecalis*, and the addition of gentamicin to daptomycin significantly improved bactericidal activity. However, the addition of rifampin decreased the activity of daptomycin against both *E. faecalis* and VRE (11). Another study indicated that the fosfomycin had activity against planktonic and adherent *E. faecalis*, and found the rifampin had no activity on planktonic or adherent *E. faecalis* (12). However, Tang HJ *et al*. found that a synergistic effect was evident using fosfomycin plus rifampin on planktonic and adherent *E. faecalis* (13). So the effective antimicrobials which work against *E. faecalis* biofilms are still little known and controversial, and there are no reports about how to treat the linezolid-resistant *E. faecalis* biofilms up to now. Thus, this study aims to explore the daptomycin, rifampin, fosfomycin alone, and daptomycin combined with rifampin or fosfomycin against the linezolid-resistant *E. faecalis* biofilms.

## MATERIALS AND METHODS

### Bacterial strains

From January 2014 to December 2016, ten linezolid-resistant *E. faecalis* isolates were collected from inpatients at 6th Affiliated Hospital of Shenzhen University Health Science Center in China. Among these linezolid-resistant isolates, four strains (FB-1, FB-2, FB-14, FB-80) which formed biofilms were used for all in vitro experiments. The strains were identified with a Phoenix 100 automated microbiology system (BD, Franklin Lakes, NJ, USA) and then two subcultured generations were re-identified with matrix-assisted laser desorption ionization time-of-flight mass spectrometry (IVD MALDI Biotyper, Germany). *E. faecalis* ATCC 29212 and OG1RF (ATCC47077) were used as reference strains. All procedures involving human participants were performed in accordance with the ethical standards of Shenzhen University and with the 1964 Helsinki declaration and its later amendments. For this type of study, formal consent is not required.

### Antimicrobial agents

Ampicillin (catalogue no. A9518), Vancomycin (catalogue no. V2002), Linezolid (catalogue no. PZ0014), Daptomycin (catalogue no. SBR00014), Rifampin (catalogue no.R3501), Gentamicin (catalogue no. E003632) and glucose-6-phosphate (catalogue no. V900924) were purchased from SIGMA-ALDRICH (Shanghai, China). Fosfomycin (catalogue no. HY-B1075) was purchased from MedChemExpress (Shanghai, China). The media were supplemented with 25 mg/liter glucose-6-phosphate for testing of fosfomycin and with 50 mg/liter Ca^2+^ for testing of daptomycin in all vitro experiments.

### Antimicrobial susceptibility testing

The MICs and the logarithmic MBC (MBClog) values for ampicillin, vancomycin, linezolid, daptomycin, rifampin and gentamicin were determined by the broth macrodilution method in cation-adjusted Mueller-Hinton broth (CAMHB), and the MICs values for fosfomycin were detected by the agar dilution method according to the Clinical and Laboratory Standards Institute guidelines (CLSI-M 1 00-S26). All experiments were performed in triplicate. The sensitivity results of antimicrobial agents were confirmed based on CLSI-M100-S26.

### Time-killing assay

The activities of daptomycin, rifampin, fosfomycin alone, and daptomycin combined with rifampin or fosfomycin (4xMIC) were determined by time-kill studies conducted with cells in the logarithmic growth phase based on the reference method (12). Briefly, the tests were performed in 14ml Polypropylene Round-Bottom Tube (FALCON 352059) in a final volume of 5 ml CAMHB and were further incubated at 37°C with shaking. At the time points of 6, 12, and 24 h, 1-ml aliquots were sampled and washed with 0.9% saline solution. Ten-fold dilutions were then plated on Muller-Hinton agar, and the numbers of CFU were determined. Medium without antimicrobial agents was used as the growth control. Bactericidal activity was defined as a ≥99.9% (i.e., ≥3-log10 CFU/ml) reduction of the initial bacterial count after 24 h, and the initial inoculum was 1.0~3.0×10^7^ CFU/ml. All experiments were performed in triplicate.

### Biofilm biomass assay

Biofilm biomasses of *E. faecalis* isolates were detected according to the reference method with minor modifications (8). Briefly, the *E. faecalis* isolates were cultivated overnight in Tryptic Soy Broth (TSB) at 37°C. Overnight cultures were diluted 1:200 in 200 μl of TSBG (TSB with 0.25% glucose) (1.0 - 3.0 × 10^7^ CFU/ml) and inoculated into 96 polystyrene microtiter plates (Costar3599, Corning). After 24 h of static incubation at 37°C (mature biofilm), the supernatant was discarded and plates were washed thrice with 0.9% saline to remove unattached cells, then the fresh TSBG containing antimicrobial agents was added to each well (200 μl/well), and the TSBG without antimicrobials was used as the growth control. After 72h of static incubation at 37°C (the medium replaced daily), the supernatant was discarded and plates were washed thrice with deionized water to remove unattached cells, stained with 1% crystal violet (CV) for 20 min at room temperature and rinsed with distilled water. Last, the CV was solubilized in ethanol-acetone (80:20, vol/vol), and optical density at 570 nm (OD_570_) was determined. The OG1RF (ATCC47077) strain was used as biofilm positive control. Each assay was performed in triplicate at least three times.

### Bacteria counting in biofilm assay

Bacteria in *E. faecalis* biofilms were determined according to the reference method (14). The *E. faecalis* isolates overnight cultures were 1:200 diluted with TSBG and inoculated into 24 polystyrene microtiter plates (1ml/well; Costar3524, Corning). After 24 h of static incubation at 37°C (mature biofilm), the supernatant was discarded and plates were washed thrice with 0.9% saline, then the fresh TSBG containing antimicrobial agents was added to each well (1ml/well), and the TSBG without antimicrobials was used as the growth control. After 72h of static incubation at 37°C (the medium replaced daily), the supernatant was discarded and plates were washed thrice with 0.9% saline, then the bacteria in the biofilms were collected by scratching the wall of the wells with a flat end toothpick and suspended in 0.9% saline. The bacteria suspension was washed twice with 0.9% saline, ten-fold diluted and then plated on Tryptic Soy agar, and the numbers of CFU were determined. All experiments were performed in triplicate.

### Detection of cell viability in mature biofilms by confocal laser scanning microscope (CLSM)

The effect of antimicrobial agents on cell viability in mature biofilms was determined using the Live/Dead Bacterial Viability method (Live/Dead BacLight, Molecular Probes, USA). The *E. faecalis* isolates overnight cultures were 1:200 diluted with TSBG and inoculated into cell-culture dishes (2ml/well; WPI, USA). After 24 h of static incubation at 37°C (mature biofilm), the supernatant was discarded and plates were washed thrice with 0.9% saline, then the fresh TSBG containing antimicrobial agents was added to each well (2ml/well), and the TSBG without antimicrobials was used as the growth control. After 72h of static incubation at 37°C (the medium replaced daily), the supernatant was discarded and plates were washed thrice with 0.9% saline. Then the mature biofilms were stained with SYTO 9 and propidium iodide (PI) at room temperature for 15 min, then observed under a Leica TCS SP8 CLSM with a 63 × 1.4-NA oil-immersion objective. Further, image analysis was performed using IMARIS 7.0.0 software (Bitplane) and the fluorescence quantities of biofilm were determined using Leica LAS AF Lite 4.0 software. All experiments were performed in triplicate and representative images were shown.

### Statistical analysis

The data were analysed using Student’s t test or nonparametric Mann–Whitney U test. *P* values <0.05 were regarded as statistically significant. All data was analyzed in SPSS software package (version 16.0, Chicago, IL,USA).

## RESULTS

### Antimicrobial susceptibility

The in vitro susceptibilities of planktonic *E. faecalis* cells were summarized in **Table 1**. All the four *E. faecalis* isolates were sensitive to ampicillin and vancomycin, but resistant to linezolid. Among these four *E. faecalis* isolates, three isolates were sensitive to daptomycin and fosfomycin, but the MIC of daptomycin to one isolate (FB-14) has reached to 8 mg/L and one isolate (FB-2) has intermediate resistance to fosfomycin. Three isolates with low level resistant to rifampin, and three isolates with the high level gentamicin MICs (≥512 mg/L) in this study.

**Table 1.** Antimicrobial susceptibility of *E.faecalis* determined by conventional broth macrodilution or agar dilution method.

| Antimicrobial agents | Susceptibility (mg/L) |  |  |  |  |  |  |  |
| --- | --- | --- | --- | --- | --- | --- | --- | --- |
|  | FB-1 |  | FB-2 |  | FB-14 |  | FB-80 |  |
|  | MIC | MBC<sub>log</sub> | MIC | MBC<sub>log</sub> | MIC | MBC<sub>log</sub> | MIC | MBC<sub>log</sub> |
| Ampicillin | 1 | 32 | 1 | 64 | 2 | 64 | 2 | 32 |
| Vancomycin | 2 | 128 | 2 | 64 | 2 | 256 | 2 | 128 |
| Linezolid | 16 | >512 | 16 | >512 | 32 | >512 | 16 | >512 |
| Daptomycin | 4 | 64 | 4 | 64 | 8 | 128 | 4 | 64 |
| Rifampin | 4 | 512 | 4 | 256 | 4 | 512 | 0.5 | 64 |
| Fosfomycin<sup>a</sup> | 64 | - | 128 | - | 64 | - | 64 | - |
| Gentamicin | >512 | >512 | >512 | >512 | 16 | 64 | 512 | >512 |

Note: MIC, minimum inhibitory concentration; MBC<sub>log</sub>, the MBC during the logarithmic growth phase; <sup>a</sup>MIC of Fosfomycin: agar dilution method;

### Antimicrobial activity on planktonic *E. faecalis* cells

The activities of daptomycin, rifampin, fosfomycin (all with 4xMIC) on planktonic *E. faecalis* cells were determined by time-killing studies. Daptomycin combined with fosfomycin or rifampin demonstrated bactericidal activities, and the daptomycin combined with fosfomycin showed better bactericidal effect than combined with rifampin on FB-1 and FB-2 isolates (**Fig. 1A and B**). Daptomycin alone, or combined with rifampin or fosfomycin indicated bactericidal activities, and daptomycin combined with fosfomycin showed best bactericidal effect on FB-14 and FB-80 isolates. Among all the four isolates, daptomycin combined with fosfomycin killed more planktonic *E. faecalis* cells (at least 2-log10 CFU/ml) than daptomycin or fosfomycin alone at the 24h of the time-kill study. It was noteworthy that rifampin or fosfomycin alone inhibited the growth of planktonic *E. faecalis* cells before 6h, but the number of bacteria increased after 6h or 12h of incubation in these four isolates (**Fig. 1**).

**Figure. 1.**
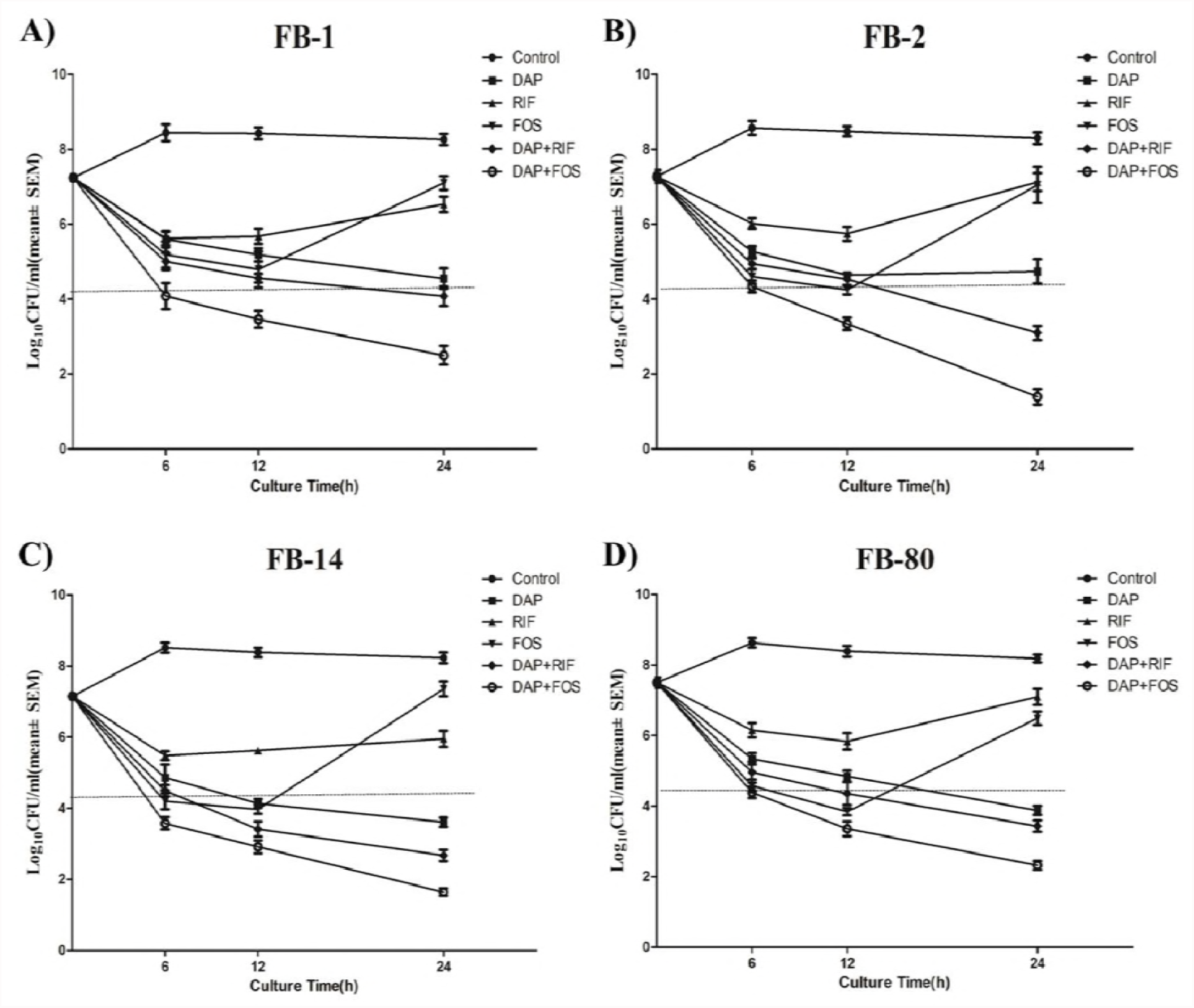
Time-kill studies for FB-1 (A), FB-2 (B), FB-14 (C), and FB-80 (D) during logarithmic growth. The horizontal dashed line represents the reduction of 3 log10 CFU/ml compared to the initial bacterial count. DAP, daptomycin; RIF, rifampin; FOS, fosfomycin. All the three agents were used at 4xMIC.

### Antimicrobial activity on the mature biofilms of *E. faecalis*

First, the activities of daptomycin, rifampin, fosfomycin (all with 16xMIC) on the mature biofilms of these four linezolid-resistant isolates were explored by microplate method with crystal violet staining. The median OD570 value was 1.45 for OG1RF (biofilm positive control strain). The daptomycin alone showed activities on the mature biofilms of FB-2, FB-14 and FB-80 isolates, and rifampin alone exhibited activity against the mature biofilms of FB-14 and FB-80 isolates (**Fig. 2**). Interestingly, daptomycin combined with fosfomycin demonstrated significantly more activity than daptomycin or fosfomycin alone against the mature biofilms of FB-2, FB-14 and FB-80 isolates. The addition of rifampin increased the activity of daptomycin against the biofilms of FB-14 and FB-80 isolates. However, daptomycin, rifampin and fosfomycin (all with 16xMIC) had no effect on the mature biofilm of FB-1 (**Fig. 2A**). Subsequently, we increased the concentrations of daptomycin from 16xMIC to 512 mg/L, and found that the high concentrations of daptomycin (512 mg/L) combined with fosfomycin also indicated good effect on the mature biofilm of FB-1 (**Fig. 3A**). Additionally, the high concentrations of daptomycin (512 mg/L) combined with fosfomycin showed more activity than 16xMIC of daptomycin combined with fosfomycin against the mature biofilms of these four linezolid-resistant isolates (**Fig. 3**).

**Figure. 2.**
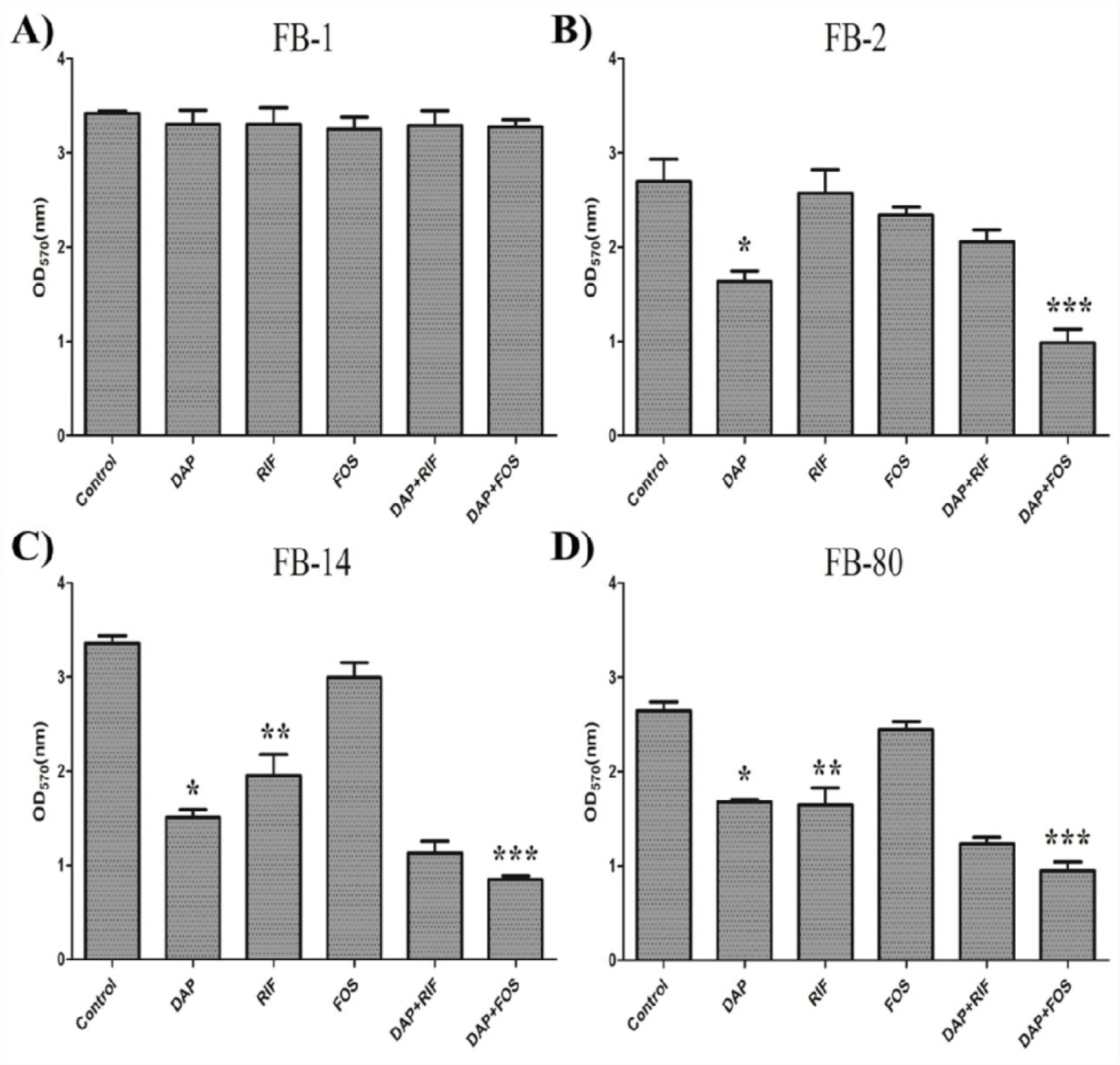
Antimicrobial activity on the mature biofilms of FB-1 (A), FB-2 (B), FB-14 (C), and FB-80 (D). The biofilm biomass of *E. faecalis* was determined by microplate method with crystal violet. DAP, daptomycin; RIF, rifampin; FOS, fosfomycin. All the three agents were used at 16xMIC. *: DAP vs Control, P<0.05; **: RIF vs Control, P<0.05; ***: DAP+FOS vs DAP or FOS alone, P<0.05.

**Figure. 3.**
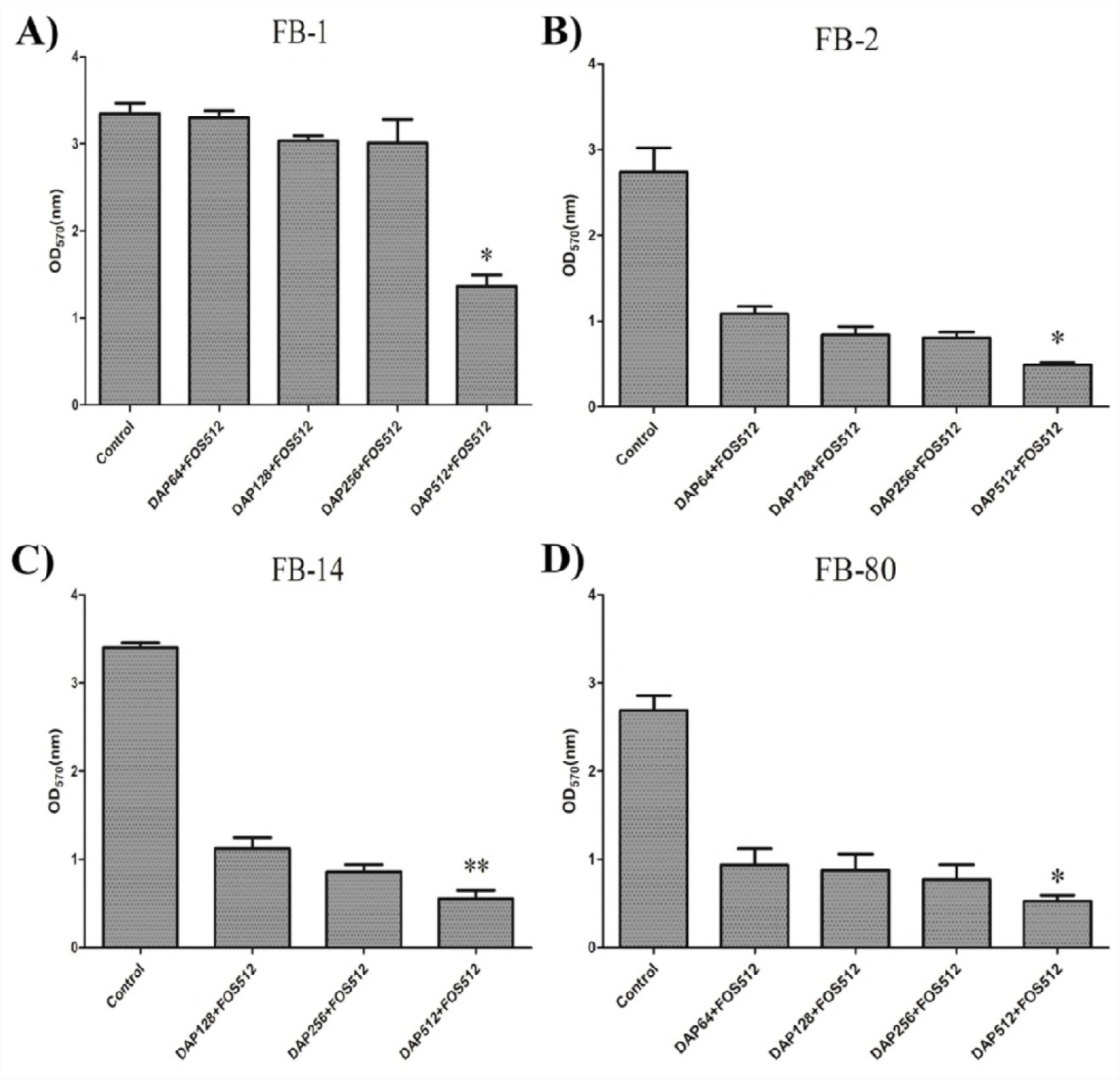
Antimicrobial activity of daptomycin combined with fosfomycin on the mature biofilms of FB-1 (A), FB-2 (B), FB-14 (C), and FB-80 (D). DAP, daptomycin, 64 (16xMIC), 128 (32xMIC), 256 (64xMIC), 512 (128xMIC) mg/L were used for FB-1, FB-2, FB-80, and 128 (16xMIC), 256 (32xMIC), 512 (64xMIC) mg/L were used for FB-14; FOS, fosfomycin, 512 mg/L were used. *: DAP512+FOS512 vs DAP64+FOS512, P<0.05; **: DAP512+FOS512 vs DAP128+FOS512, P<0.05.

### Antimicrobial agents killed the adherent cells in the mature biofilms of *E. faecalis*

How daptomycin, rifampin, and fosfomycin killed the adherent cells in the mature biofilms of *E. faecalis* were determined by the CFU numbers. First, the effects of these three agents (all with 16xMIC) on the adherent cells in the mature biofilms were detected and we found that daptomycin alone effectively killed the adherent cells in these mature biofilms of the four linezolid-resistant isolates (**Fig. 4**). Interestingly, we also found that daptomycin combined with fosfomycin killed more adherent cells than daptomycin or fosfomycin alone in these mature biofilms. The addition of rifampin also increased the activity of daptomycin against the adherent cells in the mature biofilms of FB-14 and FB-80 isolates. When the concentrations of daptomycin were increased from 16xMIC to 512 mg/L, we found that the high concentrations of daptomycin (512 mg/L) combined with fosfomycin showed significantly more killing activity than 16xMIC of daptomycin combined with fosfomycin on the adherent cells in the mature biofilms of the four linezolid-resistant isolates (**Fig. 5**).

**Figure. 4.**
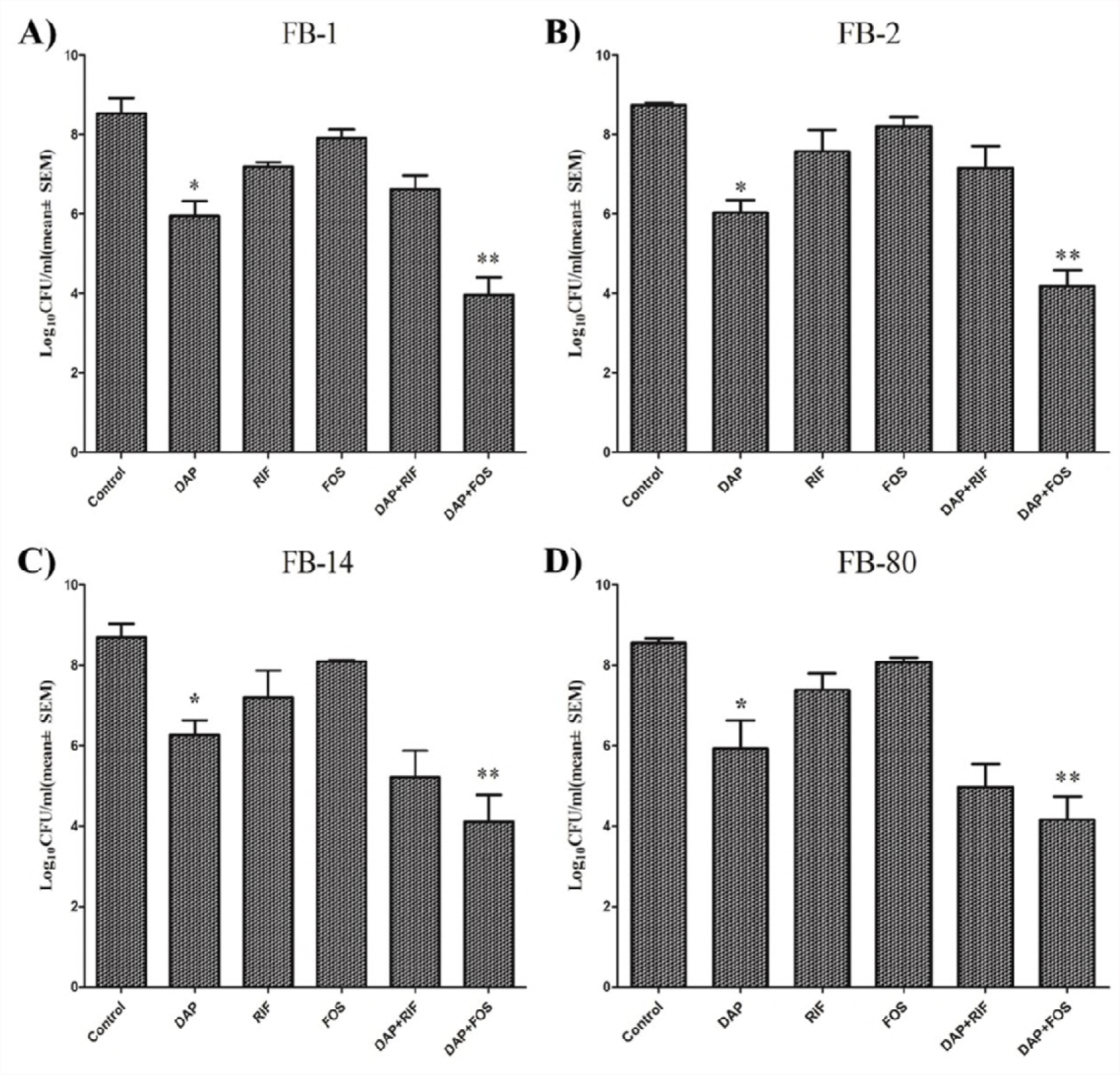
Antimicrobial agents killed the adherent cells in the mature biofilms of FB-1 (A), FB-2 (B), FB-14 (C), and FB-80 (D). The adherent cells in the mature biofilms of *E. faecalis* was determined by the CFU numbers. DAP, daptomycin; RIF, rifampin; FOS, fosfomycin. All the three agents were used at 16xMIC. *: DAP vs Control, P<0.05; **: DAP+FOS vs DAP or FOS alone, P<0.05.

**Figure. 5.**
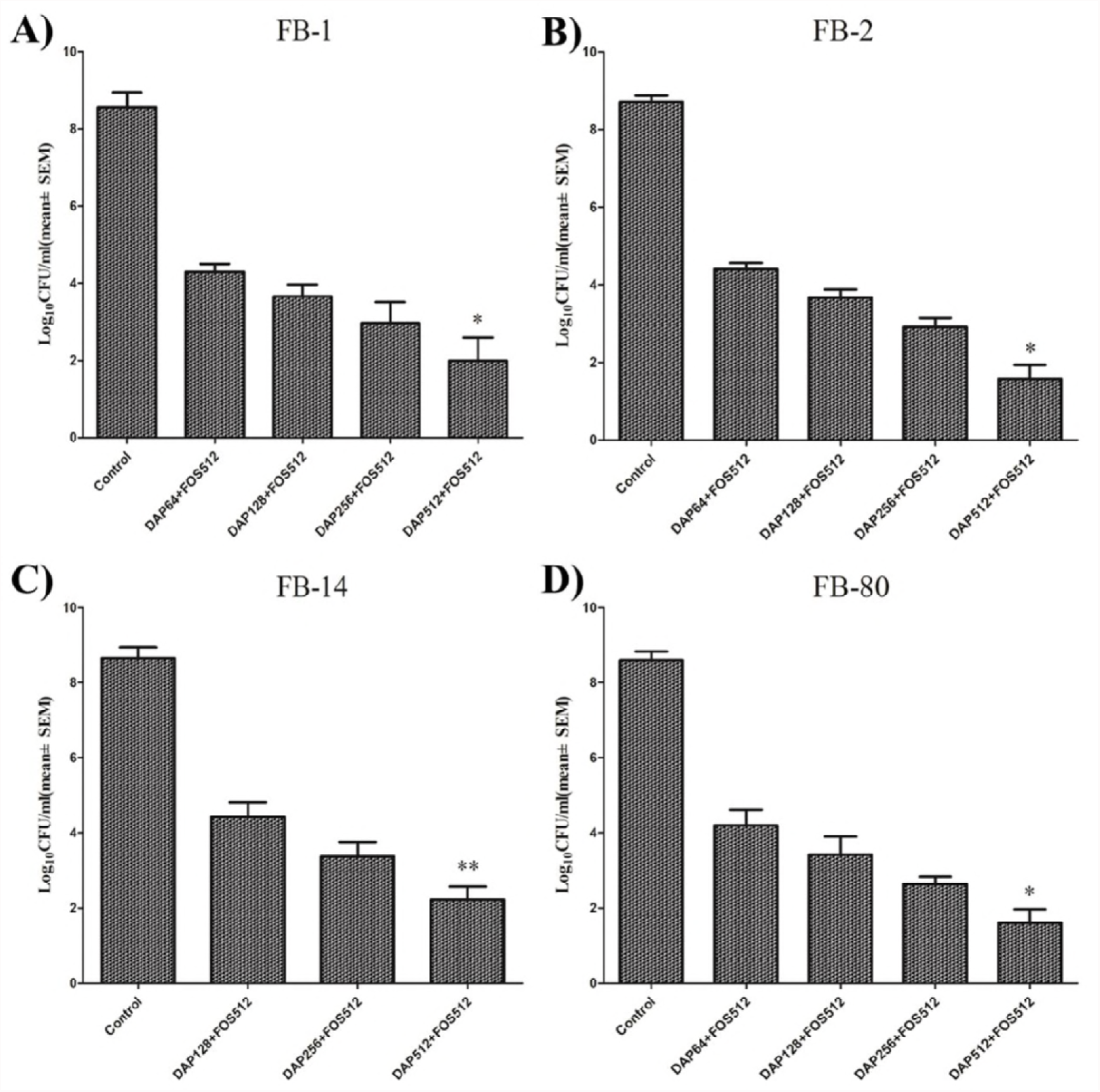
Daptomycin combined with fosfomycin killed the adherent cells in the mature biofilms of FB-1 (A), FB-2 (B), FB-14 (C), and FB-80 (D). DAP, daptomycin, 64 (16xMIC), 128 (32xMIC), 256 (64xMIC), 512 (128xMIC) mg/L were used for FB-1, FB-2, FB-80, and 128 (16xMIC), 256 (32xMIC), 512 (64xMIC) mg/L were used for FB-14; FOS, fosfomycin, 512 mg/L were used. *: DAP512+FOS512 vs DAP64+FOS512, P<0.05; **: DAP512+FOS512 vs DAP128+FOS512, P<0.05.

### Effects of daptomycin and fosfomycin on the adherent cells in the mature biofilms by CLSM

The effects of daptomycin and fosfomycin on cell viability in mature biofilms were detected by CLSM (rifampin was excepted as its red solution influenced the propidium iodide, which stained the dead cells with red fluorescence). As the **Fig. 6-9** indicated, daptomycin alone had effect on the adherent cells in these mature biofilms of the four linezolid-resistant isolates. Similar to the above results, daptomycin combined with fosfomycin showed stronger effect than daptomycin or fosfomycin alone, and the high concentrations of daptomycin (512 mg/L) combined with fosfomycin also indicated significantly stronger effect than 16xMIC of daptomycin combined with fosfomycin on the adherent cells in the mature biofilms.

**Figure. 6.**
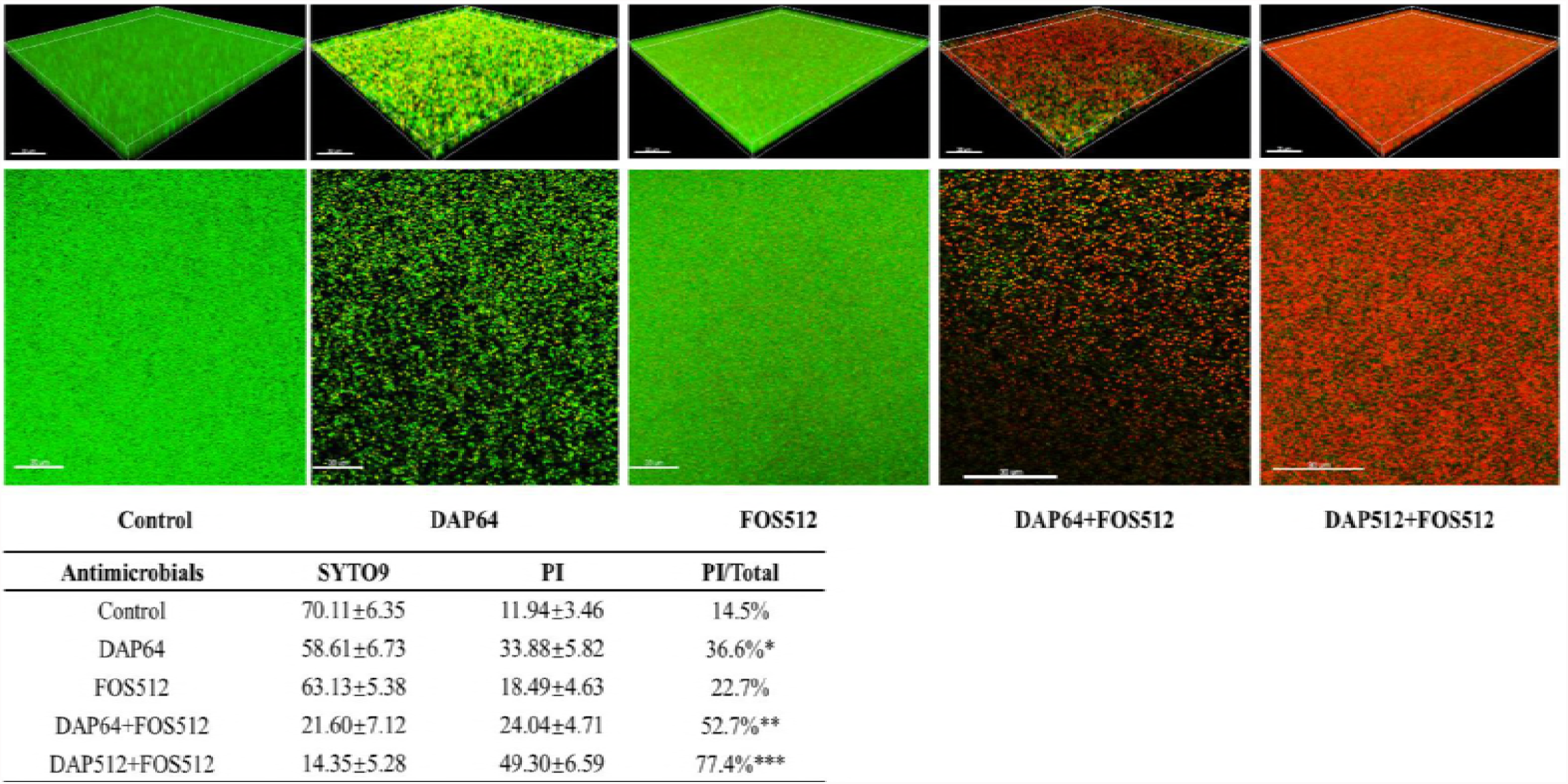
Effects of daptomycin and fosfomycin on the adherent cells in the mature biofilms of FB-1 by confocal laser scanning microscope (CLSM). The mature biofilms were stained with SYTO9 and propidium iodide (PI) and observed under a Leica TCS SP8 CLSM. Images were analyzed by IMARIS 7.0.0 software. Cells stained with green fluorescence were viable and with red fluorescence were dead. The fluorescence quantities of biofilm were determined by Leica LAS AF Lite 4.0 software. Data represent mean±SD of three independent experiments. DAP, daptomycin, 64 (16xMIC), 512 (128xMIC) mg/L were used; FOS, fosfomycin, 512 mg/L were used. *: DAP64 vs Control, P<0.05; **: DAP64+FOS512 vs DAP64 or FOS512 alone, P<0.05; ***: DAP512+FOS512 vs DAP64+FOS512, P<0.05.

**Figure. 7.**
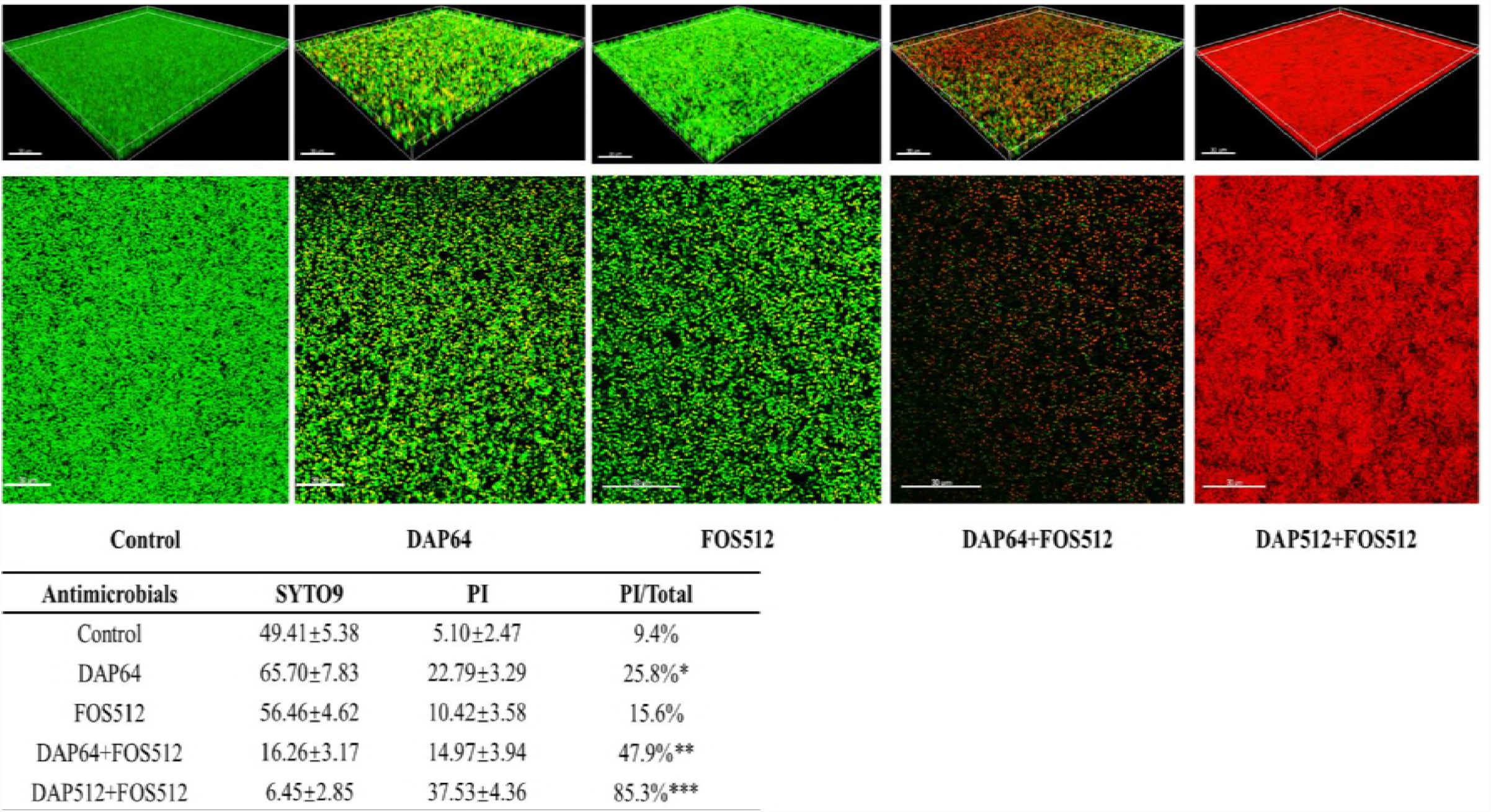
Effects of daptomycin and fosfomycin on the adherent cells in the mature biofilms of FB-2 by confocal laser scanning microscope (CLSM). The mature biofilms were stained with SYTO9 and propidium iodide (PI) and observed under a Leica TCS SP8 CLSM. Images were analyzed by IMARIS 7.0.0 software. Cells stained with green fluorescence were viable and with red fluorescence were dead. The fluorescence quantities of biofilm were determined by Leica LAS AF Lite 4.0 software. Data represent mean±SD of three independent experiments. DAP, daptomycin, 64 (16xMIC), 512 (128xMIC) mg/L were used; FOS, fosfomycin, 512 mg/L were used. *: DAP64 vs Control, P<0.05; **: DAP64+FOS512 vs DAP64 or FOS512 alone, P<0.05; ***: DAP512+FOS512 vs DAP64+FOS512, P<0.05.

**Figure. 8.**
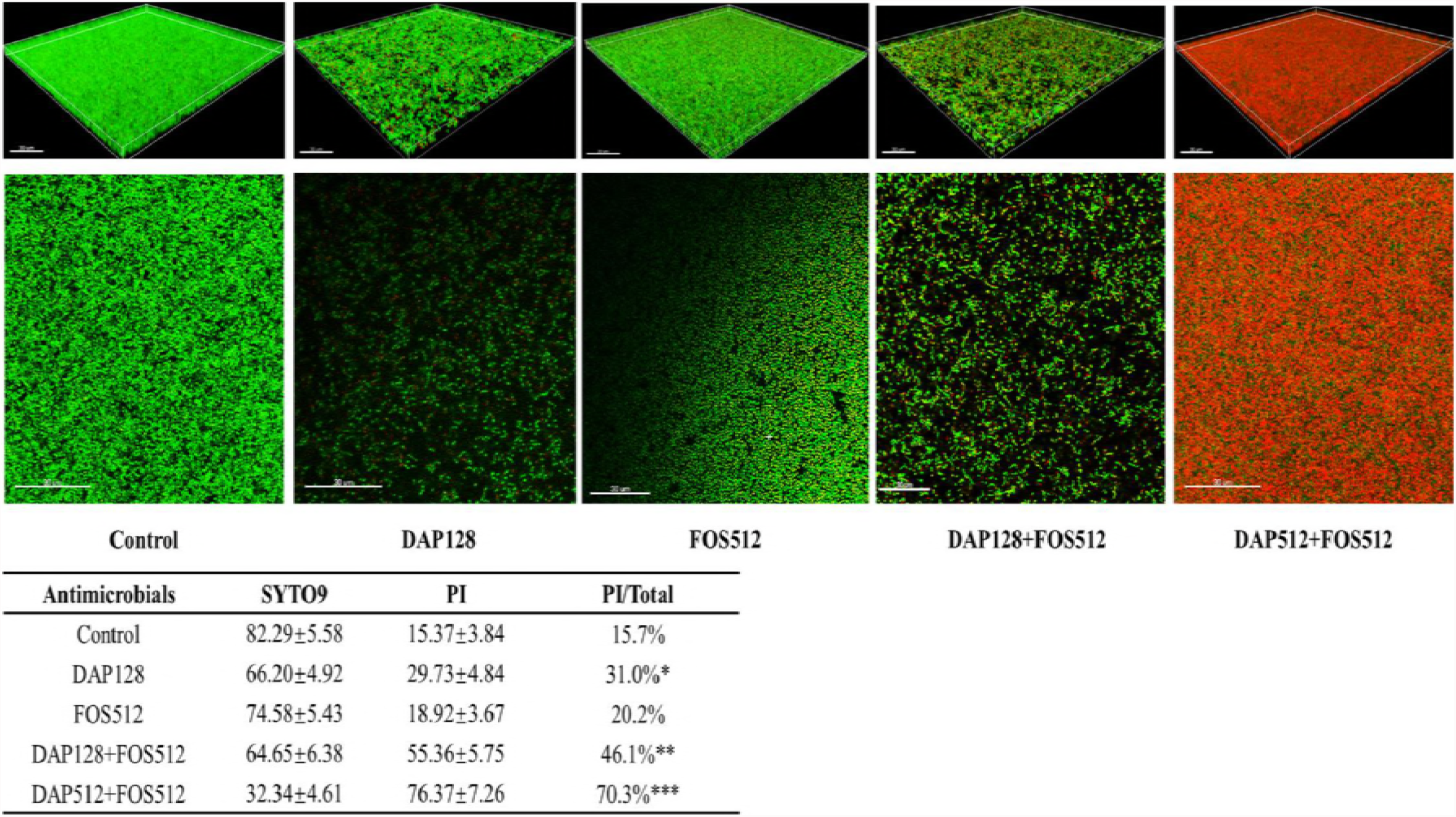
Effects of daptomycin and fosfomycin on the adherent cells in the mature biofilms of FB-14 by confocal laser scanning microscope (CLSM). The mature biofilms were stained with SYTO9 and propidium iodide (PI) and observed under a Leica TCS SP8 CLSM. Images were analyzed by IMARIS 7.0.0 software. Cells stained with green fluorescence were viable and with red fluorescence were dead. The fluorescence quantities of biofilm were determined by Leica LAS AF Lite 4.0 software. Data represent mean±SD of three independent experiments. DAP, daptomycin, 128 (16xMIC), 512 (64xMIC) mg/L were used; FOS, fosfomycin, 512 mg/L were used. *: DAP128 vs Control, P<0.05; **: DAP128+FOS512 vs DAP128 or FOS512 alone, P<0.05; ***: DAP512+FOS512 vs DAP128+FOS512, P<0.05.

**Figure. 9.**
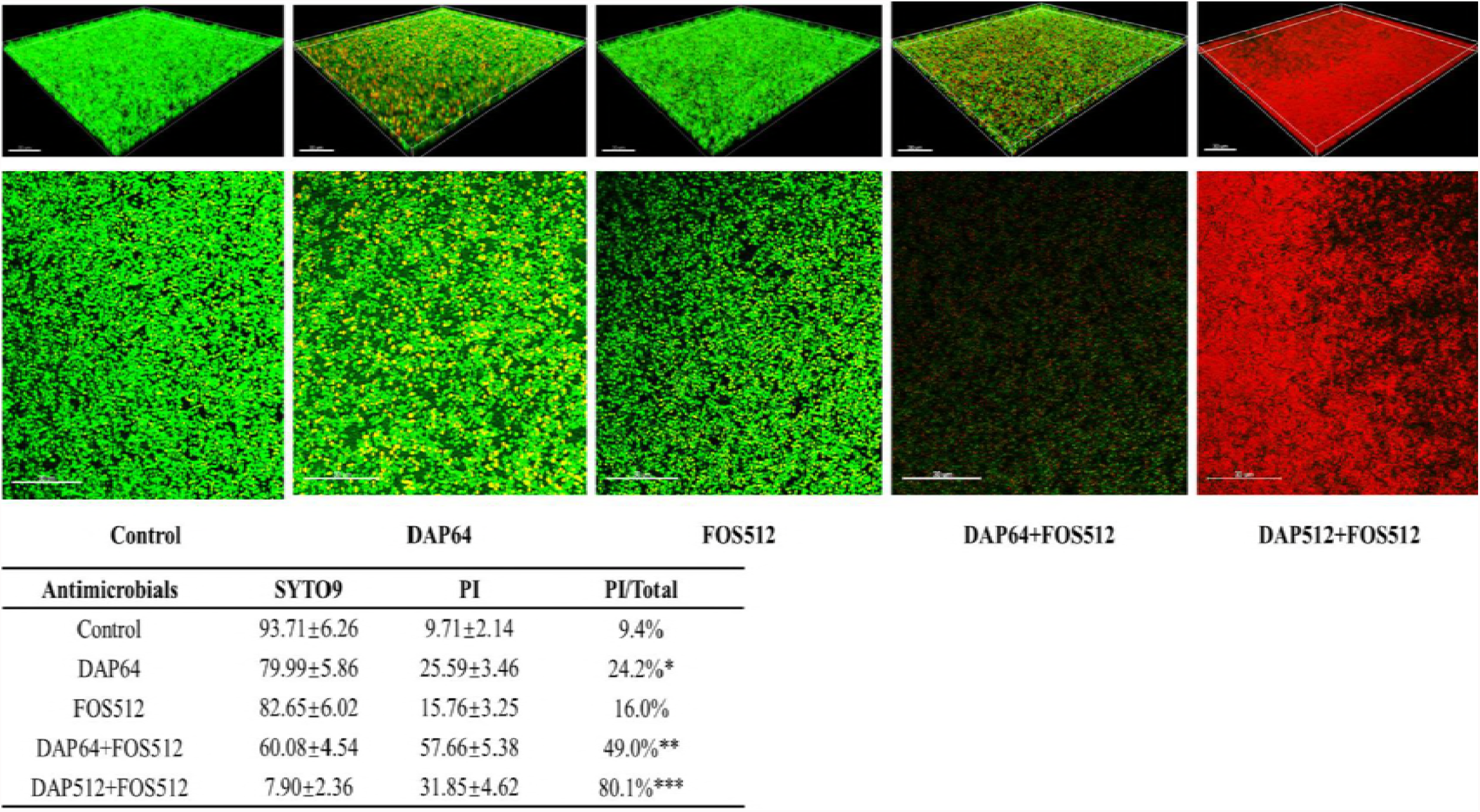
Effects of daptomycin and fosfomycin on the adherent cells in the mature biofilms of FB-80 by confocal laser scanning microscope (CLSM). The mature biofilms were stained with SYTO9 and propidium iodide (PI) and observed under a Leica TCS SP8 CLSM. Images were analyzed by IMARIS 7.0.0 software. Cells stained with green fluorescence were viable and with red fluorescence were dead. The fluorescence quantities of biofilm were determined by Leica LAS AF Lite 4.0 software. Data represent mean±SD of three independent experiments. DAP, daptomycin, 64 (16xMIC), 512 (128xMIC) mg/L were used; FOS, fosfomycin, 512 mg/L were used. *: DAP64 vs Control, P<0.05; **: DAP64+FOS512 vs DAP64 or FOS512 alone, P<0.05; ***: DAP512+FOS512 vs DAP64+FOS512, P<0.05.

## DISCUSSION

The combination of daptomycin and fosfomycin has been explored in different *E. faecalis* isolates. In the later 1980s, the combination of daptomycin and fosfomycin exhibited consistent synergistic bactericidal activity against *E. faecalis* isolates with high-level gentamicin resistance Three isolates with low level resistant to rifampin (15). Subsequently, fosfomycin was found to have synergy with daptomycin against vancomycin-resistant isolates of *E. faecium* from renal transplant patients and was also found to enhance the activity of daptomycin against vancomycin-resistant isolates of *E. faecalis* (16, 17). However, another study indicated that fosfomycin had no synergistic bactericidal effect with daptomycin on the planktonic and adherent *E. faecalis* (ATCC19433) (18). In this study, the combination of daptomycin and fosfomycin showed high bactericidal activity and a synergistic effect on the planktonic and adherent linezolid-resistant isolates of *E. faecalis*. Our results were similar with the high-level gentamicin resistant or vancomycin-resistant isolates of *E. faecalis* (15-17), but different from the *E. faecalis* ATCC19433, which was sensitive to vancomycin or linezolid (18). Why the combination of daptomycin and fosfomycin has a different effect on the linezolid-sensitive and resistant isolates of *E. faecalis* is still unknown and needs further exploration.

The high-dose daptomycin (10 mg/kg/day) plus fosfomycin has been proven to be safe and effective in treating *S. aureus* endocarditis (19). Another study also found that the high-dose daptomycin (≥8 mg/kg/day) was effective and safe for the treatment of infective endocarditis, which is mostly caused by methicillin-resistant *S. aureus* and vancomycin-resistant *E. faecium* (20). Similar to the above research, the present study also showed that the high concentrations of daptomycin (512 mg/L) combined with fosfomycin had a significantly stronger effect on the mature biofilms and the adherent cells of the linezolid-resistant isolates than 16xMIC of daptomycin combined with fosfomycin. Thus, the patients infected with linezolid-resistant *E. faecalis* may also benefit from treatment with high-dose daptomycin, but this issue needs further in vivo studies.

Fosfomycin has indicated activity against both Gram-positive and Gram-negative biofilms, such as *pseudomonas aeruginosa*, *Escherichia coli* and *S. aureus* (21-23). There are several studies that have reported fosfomycin is effective against *E. faecalis* biofilms, but this is still controversial. Oliva A *et al* indicated that fosfomycin alone cleared planktonic bacteria from 74% of cage fluids and eradicated biofilm bacteria from 43% of cages in their study (12). However, another study found that among vancomycin-resistant *E. faecalis* and *E. faecium* isolates, fosfomycin alone had no bactericidal effect on the planktonic and adherent bacteria (13). Our study also showed that fosfomycin alone had no significantly killing activity on the planktonic and adherent cells among these linezolid-resistant isolates of *E. faecalis.* Thus, the role of fosfomycin in *E. faecalis* biofilm infections has not been widely investigated and needs further confirmation. In addition to the uncertainty of the effect of fosfomycin alone on the biofilm-related infections, prolonged therapy with fosfomycin may promote the emergence of fosfomycin-resistant isolates (24). So fosfomycin is not recommended for monotherapy in clinical practice, and fosfomycin-included combined treatment may provide better options in these biofilm-related infections.

Rifampin alone or combined with linezolid or vancomycin achieved good effects on the biofilms of methicillin-resistant *Staphylococcus aureus* (MRSA) strains, and was even effective against the implant-associated infections which are caused by MRSA (25-27). However, the role of rifampin in enterococcal infection remains confusing and controversial. Rifampin was explored effectively against the biofilms of vancomycin sensitive *E. faecalis* in combination with ciprofloxacin and linezolid in vitro, and in combination with tigecycline in vivo studies (28, 29). Tang HJ *et al*. also found that rifampin combined with fosfomycin indicated a synergistic effect on the planktonic and adherent *E. faecalis* isolates, which are resistant to vancomycin (13). In contrast to the above results, Oliva A *et al*. found rifampin with no activity against enterococcal biofilms of *E. faecalis* ATCC19433 (sensitive to vancomycin), either in vitro or in vivo (12). Another study showed the addition of rifampin even decreased the activity of daptomycin against the biofilms of vancomycin-susceptible *E. faecalis* (11). However, the present study found the addition of rifampin increased the activity of daptomycin against the planktonic *E. faecalis* isolates, which are more obvious in FB-2 and FB-14 strains. This study also found that rifampin increased the activity of daptomycin on the adherent cells and mature biofilms among FB-14 and FB-80 isolates, but was not observed in FB-1 and FB-2 strains. Based on the previous and present studies, rifampin indicated disparate effects on the different *E. faecalis* isolates, and was not related to the antimicrobial susceptibility, such as vancomycin or linezolid.

In conclusion, this study indicated that daptomycin combined with fosfomycin works effectively against the planktonic and adherent linezolid-resistant isolates of *E. faecalis.* The high concentrations of daptomycin combined with fosfomycin achieved significantly stronger effect on these isolates. However, the role of rifampin in these linezolid-resistant isolates of *E. faecalis* is inconsistent and needs more studies to resolve this issue.

## ACKNOWLEDGMENTS

The authors thank Prof. Xiao-gang Xu (Institute of Antibiotics, Huashan Hospital, Fudan University, Shanghai, China) for his suggestions on the experiment schemes and thank MS. Cynthia Brast (University of Florida, Orlando, USA) for her review on this manuscript.

## FUNDING

This work was supported by grants from the National Natural Science Foundation of China (grant number 81170370); the Sanming Project of Medicine in Shenzhen (grant number SMGC201705029); the Shenzhen Scientific Research Program (grant numbers JCYJ20170412143551332, JCYJ20170307153714512, JCYJ20170307153425389, JCYJ20170307153919735); the Scientific Research Project of the Shenzhen Health and Family Planning System (grant number 201601058); and the Shenzhen Nanshan District Scientific Research Program of the People’s Republic of China (grant number 2016010).

## CONFLICT of INTEREST

The authors declare that they have no conflicts of interest.

## REFERENCES

1. Beganovic M, Luther MK, Rice LB, Arias CA, Rybak MJ, LaPlante KL. 2018. A Review of Combination Antimicrobial Therapy for *Enterococcus Faecalis* Bloodstream Infections and Infective Endocarditis. Clin Infect Dis. https://doi:10.1093/cid/ciy064. [Epub ahead of print]

2. Flokas ME, Karageorgos SA, Detsis M, Alevizakos M, Mylonakis E. 2017. Vancomycin-resistant enterococci colonisation, risk factors and risk for infection among hospitalised paediatric patients: a systematic review and meta-analysis. Int J Antimicrob Agents 49: 565–572. https://doi:10.1016/j.ijantimicag.2017.01.008.

3. Lazaris A, Coleman DC, Kearns AM, Pichon B, Kinnevey PM, Earls MR, et al. 2017. Novel multiresistance cfr plasmids in linezolid-resistant methicillin-resistant *Staphylococcus epidermidis* and vancomycin-resistant *Enterococcus faecium* (VRE) from a hospital outbreak: co-location of cfr and optrA in VRE. J Antimicrob Chemother 72: 3252–3257. https://doi:10.1093/jac/dkx292.

4. Sandoe J A, Witherden IR, Cove JH, Heritage J, Wilcox MH. 2003. Correlation between enterococcal biofilm formation in vitro and medical-device-related infection potential in vivo. J Med Microbiol 52: 547–550.

5. Seno Y, Kariyama R, Mitsuhata R, Monden K, Kumon H. 2005. Clinical implications of biofilm formation by *Enterococcus faecalis* in the urinary tract. Acta Med Okayama 59: 79–87.

6. Toledo-Arana A, Valle J, Solano C, Arrizubieta M, Cucarella C, Lamata M, et al. 2001. The enterococcal surface protein, Esp, is involved in *Enterococcus faecalis* biofilm formation. Appl Environ Microbiol 67: 4538–4545.

7. Dupre I, Zanetti S, Schito AM, Fadda G, Sechi LA. 2003. Incidence of virulence determinants in clinical *Enterococcus faecium* and *Enterococcus faecalis* isolates collected in Sardinia (Italy). J Med Microbiol 52: 491–498.

8. Mohamed JA, Huang W, Nallapareddy SR, Teng F, Murray BE. 2004. Influence of origin of isolates, specially endocarditis isolates, and various genes on biofilm formation by *Enterococcus faecalis*. Infect Immun 72: 3658–3663.

9. Zheng JX, Wu Y, Lin ZW, Pu ZY, Yao WM, Chen Z, et al. 2017. Characteristics of and Virulence Factors Associated with Biofilm Formation in Clinical *Enterococcus faecalis* Isolates in China. Front Microbiol 8: 2338. https://doi:10.3389/fmicb.2017.02338.

10. Lewis K. Riddle of biofilm resistance. 2001. Antimicrob Agents Chemother 45: 999–1007.

11. Luther MK, Arvanitis M, Mylonakis E, LaPlante KL. 2014. Activity of Daptomycin or Linezolid in Combination with Rifampin or Gentamicin against Biofilm-Forming *Enterococcus faecalis* or *E. faecium* in an In Vitro Pharmacodynamic Model Using Simulated Endocardial Vegetations and an In Vivo Survival Assay Using Galleria mellonella Larvae. Antimicrob Agents Chemother 58: 4612–4620. https://doi:10.1128/AAC.02790-13.

12. Oliva A, Furustrand Tafin U, Maiolo EM, Jeddari S, Bétrisey B, Trampuz A. 2014. Activities of Fosfomycin and Rifampin on Planktonic and Adherent *Enterococcus faecalis* Strains in an Experimental Foreign-Body Infection Model. Antimicrob Agents Chemother 58: 1284–1293. https://doi:10.1128/AAC.02583-12.

13. Tang HJ, Chen CC, Zhang CC, Su BA, Li CM, Weng TC, et al. 2013. In vitro efficacy of fosfomycin-based combinations against clinical vancomycin-resistant *Enterococcus* isolates. Diagn Microbiol Infect Dis 77: 254–257. https://doi:10.1016/j.diagmicrobio.2013.07.012.

14. Wang C, Li M, Dong D, Wang J, Ren J, Otto M, et al. 2007. Role of ClpP in biofilm formation and virulence of *Staphylococcus epidermidis*. Microbes Infect 9: 1376–1383.

15. Rice LB, Eliopoulos GM, Moellering RC Jr. 1989. In Vitro Synergism between Daptomycin and Fosfomycin against *Enterococcus faecalis* Isolates with High-Level Gentamicin Resistance. Antimicrob Agents Chemother 33:470–473.

16. Descourouez JL, Jorgenson MR, Wergin JE, Rose WE. 2013. Fosfomycin Synergy In Vitro with Amoxicillin, Daptomycin, and Linezolid against Vancomycin-Resistant *Enterococcus faecium* from Renal Transplant Patients with Infected Urinary Stents. Antimicrob Agents Chemother 57:1518–1520. https://doi:10.1128/AAC.02099-12.

17. Hall Snyder AD, Werth BJ, Nonejuie P, McRoberts JP, Pogliano J, Sakoulas G, et al. 2016. Fosfomycin Enhances the Activity of Daptomycin against Vancomycin-Resistant Enterococci in an In Vitro Pharmacokinetic-Pharmacodynamic Model. Antimicrob Agents Chemother 60:5716–5723. https://doi:10.1128/AAC.00687-16.

18. Oliva A, Furustrand Tafin U, Maiolo EM, Jeddari S, Bétrisey B, Trampuz A. 2014. Activities of Fosfomycin and Rifampin on Planktonic and Adherent *Enterococcus faecalis* Strains in an Experimental Foreign-Body Infection Model. Antimicrob Agents Chemother 58:1284–1293. https://doi:10.1128/AAC.02583-12.

19. Miró JM, Entenza JM, Del Río A, Velasco M, Castañeda X, Garcia de la Mària C, et al. 2012. High-Dose Daptomycin plus Fosfomycin Is Safe and Effective in Treating Methicillin-Susceptible and Methicillin-Resistant *Staphylococcus aureus* Endocarditis. Antimicrob Agents Chemother 56: 4511–4515.

20. Kullar R, Casapao AM, Davis SL, Levine DP, Zhao JJ, Crank CW, et al. 2013. A multicentre evaluation of the effectiveness and safety of high-dose daptomycin for the treatment of infective endocarditis. J Antimicrob Chemother 68: 2921–2926. https://doi:10.1093/jac/dkt294.

21. Mikuniya T, Kato Y, Ida T, Maebashi K, Monden K, Kariyama R, et al. 2007. Treatment of *Pseudomonas aeruginosa* biofilms with a combination of fluoroquinolones and fosfomycin in a rat urinary tract infection model. J Infect Chemother 13: 285–290.

22. Corvec S, Furustrand Tafin U, Betrisey B, Borens O, Trampuz A. 2013. Activities of fosfomycin, tigecycline, colistin, and gentamicin against extended-spectrum lactamase-producing *Escherichia coli* in a foreignbody infection model. Antimicrob Agents Chemother 57:1421–1427.

23. Garrigos C, Murillo O, Lora-Tamayo J, Verdaguer R, Tubau F, Cabellos C, et al. 2013. Fosfomycin-daptomycin and other fosfomycin combinations as alternative therapies in experimental foreign-body infection by methicillin-resistant *Staphylococcus aureus*. Antimicrob Agents Chemother 57: 606–610. https://doi:10.1128/AAC.01570-12.

24. Raz R. 2012. Fosfomycin: an old-new antibiotic. Clin Microbiol Infect 18: 4–7. https://doi:10.1111/j.1469-0691.2011.03636.x.

25. Zimmerli W, Widmer AF, Blatter M, Frei R, Ochsner PE. 1998. Role of rifampin for treatment of orthopedic implant-related staphylococcal infections: a randomized controlled trial. Foreign-Body Infection (FBI) Study Group. JAMA 279: 1537–1541.

26. Baldoni D, Haschke M, Rajacic Z, Zimmerli W, Trampuz A. 2009. Linezolid alone or combined with rifampin against methicillin-resistant *Staphylococcus aureus* in experimental foreign-body infection. Antimicrob Agents Chemother 53:1142–1148. https://doi:10.1128/AAC.00775-08.

27. Vergidis P, Rouse MS, Euba G, Karau MJ, Schmidt SM, Mandrekar JN, et al. 2011.Treatment with linezolid or vancomycin in combination with rifampin is effective in an animal model of methicillin-resistant *Staphylococcus aureus* foreign body osteomyelitis. Antimicrob Agents Chemother 55:1182–1186. https://doi:10.1128/AAC.00740-10.

28. Holmberg A, Morgelin M, Rasmussen M. 2012. Effectiveness of ciprofloxacin or linezolid in combination with rifampicin against *Enterococcus faecalis* in biofilms. J Antimicrob Chemother 67:433–439. https://doi:10.1093/jac/dkr477.

29. Minardi D, Cirioni O, Ghiselli R, Silvestri C, Mocchegiani F, Gabrielli E, et al. 2012. Efficacy of tigecycline and rifampin alone and in combination against *Enterococcus faecalis* biofilm infection in a rat model of ureteral stent. J Surg Res 176: 1–6. https://doi:10.1016/j.jss.2011.05.002.

